# Transmembrane Stem Cell Factor Protein Therapeutics Enhance Revascularization in Ischemia without Mast Cell Activation

**DOI:** 10.1101/2020.04.06.028563

**Authors:** Eri Takematsu, Jeff Auster, Po-Chih Chen, Sanjana Srinath, Sophia Canga, Aditya Singh, Marjan Majid, Michael Sherman, Andrew Dunn, Annette Graham, Patricia Martin, Aaron B. Baker

**Affiliations:** Department of Biomedical Engineering, University of Texas at Austin, Austin, TX; Department of Biochemistry & Molecular Biology, University of Texas Medical Branch, Galveston, TX; Department of Biological and Biomedical Sciences, School of Health and Life Sciences, Glasgow Caledonian University, G4 0BA, Scotland, UK; Institute for Cellular and Molecular Biology, University of Texas at Austin, Austin, TX; The Institute for Computational Engineering and Sciences, University of Texas at Austin, Austin, TX; Institute for Biomaterials, Drug Delivery and Regenerative Medicine, University of Texas at Austin, Austin, TX

**Keywords:** stem cell factor, transmembrane stem cell factor, nanodiscs, ischemia, therapeutic angiogenesis, revascularization, peripheral vascular disease

## Abstract

Stem cell factor (SCF) is a cytokine that regulates hematopoiesis and other biological processes. While clinical treatments using SCF would be highly beneficial, these have been limited by toxicity related to mast cell activation. Transmembrane SCF (tmSCF) has differential activity from soluble SCF and has not been explored as a therapeutic agent. We created novel therapeutics using tmSCF embedded in proteoliposomes or lipid nanodiscs. Mouse models of anaphylaxis and ischemia revealed the tmSCF-based therapies did not activate mast cells and improved the revascularization in the ischemic hind limb. Proteoliposomal tmSCF preferentially acted on endothelial cells to induce angiogenesis while tmSCF nanodiscs had greater activity in inducing stem cell mobilization and recruitment to the site of injury. The type of lipid nanocarrier used altered the relative cellular uptake pathways and signaling in a cell type dependent manner. Overall, we found that tmSCF-based therapies can provide therapeutic benefits without off target effects.

## Introduction

Over the past three decades, protein therapeutics have emerged as a powerful approach to drug development.^1,2^ The first use of a therapeutic protein developed was insulin, used as a therapy for diabetes mellitus.^3^ Since then, over 200 protein-based compounds have been approved for clinical use and over 250 proteins are currently in various stages of clinical evaluation.^4,5^ While proteins have immense potential in bridging the gap between a scientific discovery and translation into clinical therapy, there remain major limitations and challenges in their delivery and efficacy.^6^ This has been particularly the case with membrane proteins as these molecules are natively found in the complex lipid bilayer of the cell membrane and often require this environment for proper function and solubility.^7^ Virtually all disease processes involve membrane proteins as receptors, co-receptors or membrane bound factors that are needed to transmit cellular signals.^8–10^ Thus, the delivery of membrane protein therapeutics may provide a rich strategy for enabling next generation of protein therapeutics.

Stem cell factor (SCF) is a hematopoietic cytokine that signals through the c-Kit receptor (CD117)^11^, and is also known as Kit ligand, Steel factor or mast cell growth factor. Through alternative splicing SCF is expressed in cells initially as a transmembrane protein that subsequently enzymatically cleaved into soluble SCF or a shorter isoform that lacks the cleavable domain and remains membrane bound as transmembrane SCF (tmSCF)^12^. Signaling through SCF induced c-Kit activation is key to the maintenance of the hematopoietic stem cells (HSCs) and progenitor cells in the bone marrow^13,14^. There are several potential uses for SCF in therapeutic applications including improving the survival and expansion of HSCs following exposure to radiation^15,16^, inducing neuroprotective effects following stroke^17–19^, and enhancing the recovery of the heart following myocardial infarction. However, SCF also has an important role in regulating mast cell maturation and activation^20,21^. Treatment with exogenous SCF leads to mast cell activation and anaphylaxis in animal studies and clinical trials, severely limiting its therapeutic application^22–29^.

In the body, SCF is expressed both as a longer isoform that is initially a transmembrane protein but then released as soluble SCF through enzymatic cleavage and as a shorter isoform that remains as transmembrane SCF^30^. Transmembrane SCF is found in the stromal cells of the bone marrow where it functions to support the proliferation and survival of progenitor cells^31^. Soluble and transmembrane SCF differ in terms of their ability to activate the c-Kit receptor, induce cellular responses and ability to promote adhesion between hematopoietic stem cells and extracellular matrix^11,32^. Several reports have further demonstrated that tmSCF can induce prolonged activation of the c-Kit receptor^33^ and longer term proliferation of CD34^+^ hematopoietic cells in comparison to soluble SCF^34^. While studies have supported that gene expression of tmSCF can improve recovery following myocardial infarction^22^, delivery of exogenous tmSCF has not been explored as therapeutic strategy in preclinical or clinical studies.

In this study, we developed therapeutics based on tmSCF delivered in lipid nanodiscs or as proteoliposomes. Our studies demonstrated that tmSCF-based therapies do not induce mast cell activation in mice, in contrast to soluble SCF. Further, we found that tmSCF nanodiscs and proteoliposomes enhanced the recovery from hind limb ischemia of both wild type and ob/ob mice. Trafficking studies demonstrated that mast cells mainly use clathrin-mediated pathways to uptake SCF, while they use both clathrin- and caveolin-mediated pathways to internalize tmSCF therapies. In addition, the type of nanocarrier used altered the signaling, internalization pathways, and relative activity towards in endothelial cells or endothelial progenitor cells. Overall, our studies suggest that therapeutics based on tmSCF can enhance stem cell recruitment and revascularization without toxic activities including mast cell activation.

## Results

### Synthesis of transmembrane stem cell factor nanodiscs and proteoliposomes

We created several formulations of recombinant tmSCF to test the effectiveness and differential response in comparison to soluble SCF. Our previous studies had identified that embedding transmembrane proteins in proteoliposomes can alter their therapeutic and signaling properties^35,36^. In addition, recent work has identified methods for creating lipid nanodiscs that are stabilized by membrane scaffolding proteins (MSPs). Thus, we aim to create two types of nanocarriers, tmSCF proteoliposomes and nanodiscs to deliver tmSCF effectively to ischemic sites (**Fig. 1E**). Transmembrane stem cell factor (tmSCF) protein was first harvested and purified as described in method section. SDS-PAGE, western blotting and silver staining were performed to analyze the purity of the final concentrated samples. The results of silver staining and western blotting confirmed the purity of tmSCF protein (**Supplemental Fig. 1**). We measured the size of the purified tmSCF and found that there is likely self-association/aggregation in the absence of a lipid carrier (**Supplemental Fig. 2**). We fabricated tmSCF proteoliposomes (tmSCFPLs) using detergent depletion as previously described^35,36^. The initial liposomes were around 350 nm in diameter as expected from the extrusion membrane used (**Fig. 1F**). After embedding tmSCF, the size was increased to 450 nm. We further verified the liposomal structure using TEM and cryo-EM (**Fig. 1G**). Another type of nanocarrier is nanodiscs made by 1-palmitoyl-2-oleoyl-sn-glycero-3-phosphocholine (POPC) lipid and Membrane Scaffold Protein 1D1 (MSP1D1). MSP1D1 comes together to form a “lariat” structure that stabilizes the hydrophobic edges of a lipid nanodisc. Purified nandiscs had a size of 20-30 nm and following tmSCF embedding and detergent depletion, the size increased to 150 nm (**Fig. 1H**). The structure of the nanodiscs was further verified using TEM and cryo-EM, which revealed nanodisc structures (**Fig. 1G**).

**Figure 1.**
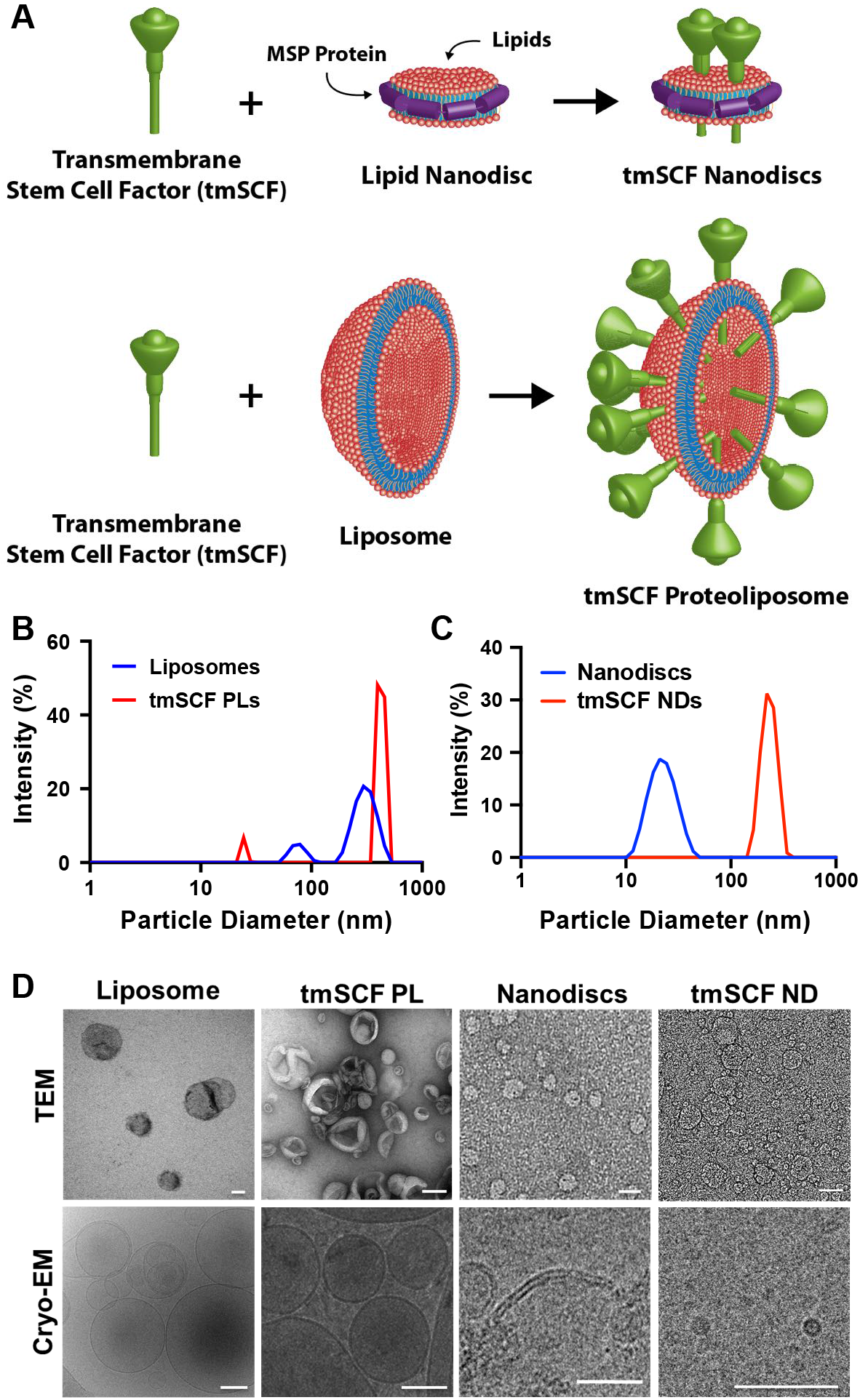
SCF is deficient in the skin of patients with diabetes. Immunostaining was conducted on the skin of non-diabetic (ND) and type 2 diabetic patients (T2D). (A) Schematic illustration of tmSCF proteoliposomes (tmSCFPLs) and tmSCF nanodiscs (tmSCFNDs). (B) Size distribution for liposomes and proteoliposome with tmSCF measured by dynamic light scattering. (C) Size distribution for nanodiscs and tmSCF nanodiscs. (D) Representative TEM and cryo-EM images of liposomes, tmSCFPLs, nanodiscs, and tmSCFNDs. Scale bar = 100 nm.

**Figure 2.**
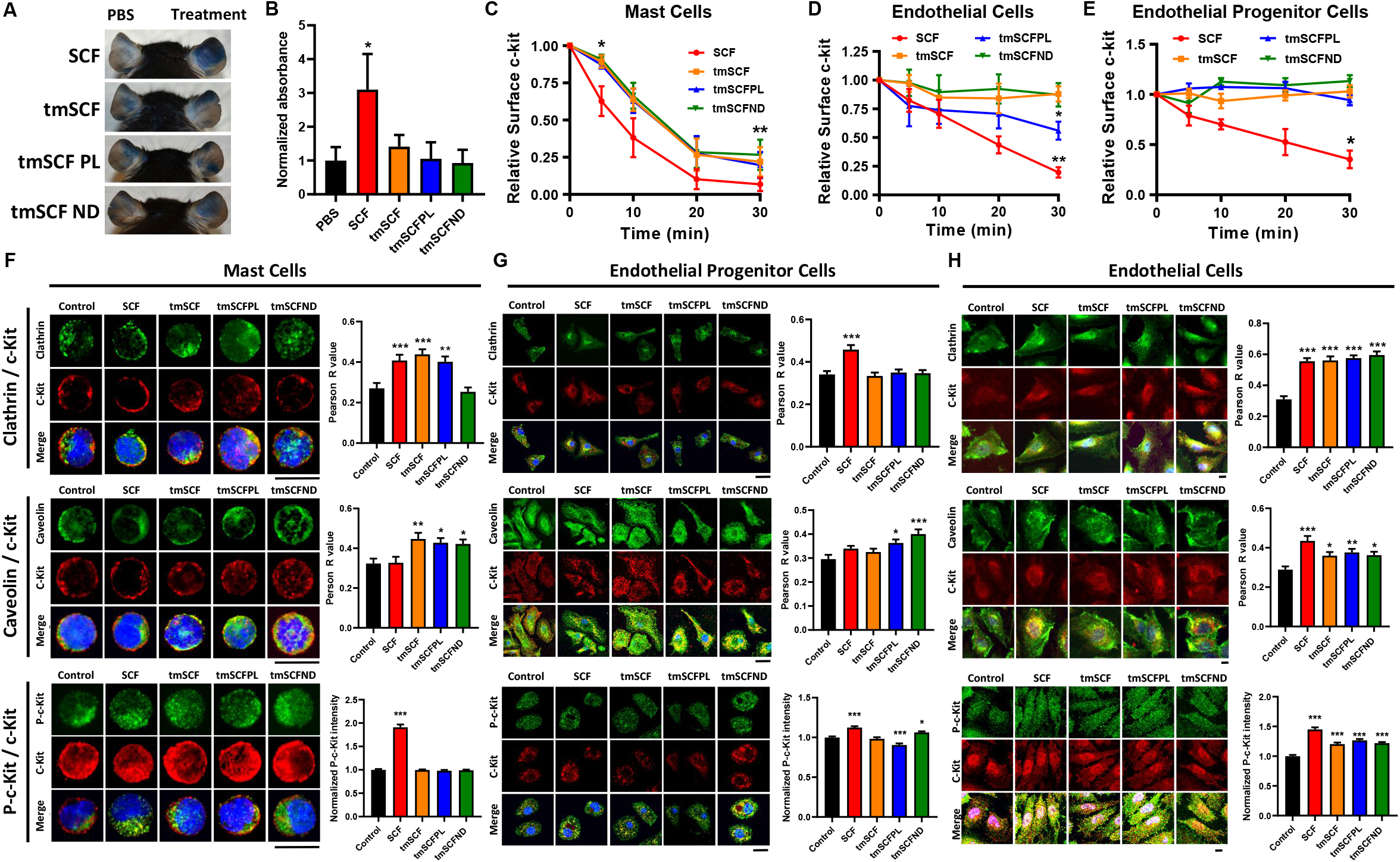
Nanocarriers serves as a switch to control protein uptake mechanism. (A) Representative pictures of mice ears after Evan’s blue extravasation assay. PBS was injected to the left ear as a control and treatments were injected to the right ear. (B) Quantification result of the absorbance at 620 nm (n = 7-8). (C) Surface staining for c-Kit on MC9 mast cells was monitored by flow cytometry. The intensity was normalized to 0 min time point to evaluate the treatment uptake kinetics (n = 6). (D) Surface c-Kit on bone marrow mononuclear cells were monitored by flow cytometry (n = 3-6). (E) Surface c-Kit on HUVECs were monitored by flow cytometry (n = 4-8). (F) (Top) Representative pictures of single mast cell stained with clathrin and c-Kit. Pearson’s R value of the colocalization of clathrin and c-Kit (n = 30). (Middle) Representative pictures of single mast cell stained with caveolin and c-Kit. Pearson’s R value of the colocalization of caveolin and c-Kit (n = 30). (Bottom) Representative pictures of single mast cell stained with c-Kit and p-C-kit. P-c-Kit mean intensity inside of mast cells was quantified using photoshop (n = 30). (G) (Top) Representative pictures of EPCs stained with clathrin and c-Kit. Pearson’s R value of the colocalization of clathrin and c-Kit (n = 30). (Middle) Representative pictures of EPCs stained with caveolin and c-Kit. Pearson’s R value of the colocalization of caveolin and c-Kit (n = 30). Representative pictures of EPCs stained with c-Kit and p-c-Kit. P-c-Kit mean intensity inside of EPCs were quantified using photoshop (n = 30). (H) (Top) Representative pictures of HUVECs stained with clathrin and c-Kit. Pearson’s R value of the colocalization of clathrin and c-Kit (n = 30). (Middle) Representative pictures of HUVECs stained with caveolin and c-Kit. Pearson’s R value of the colocalization of caveolin and c-Kit (n = 30). (Bottom) Representative pictures of HUVECs stained with c-Kit and p-c-Kit. P-c-Kit mean intensity inside of HUVECs were quantified using photoshop (n = 30). Scale bar is 30 μm. *p<0.05 versus control, **p<0.01 versus control, ***p<0.001 versus control.

### Transmembrane stem cell factor-based treatments do not induce mast cell activation

A major limitation in the clinical use of soluble SCF is the activation of mast cells and potential induction of anaphylaxis^21,37^. To test whether tmSCF activates mast cells, we injected mice with Evan’s blue dye and then locally injected PBS, soluble SCF, tmSCF, tmSCFPLs or tmSCFNDs into the ears of the mice. After three hours, we imaged the ears and extracted the dye with DMSO to quantify the vascular leakage induced by the treatments. We found that SCF induced vascular leakage but tmSCF-based therapies did not induce this response (**Fig. 2A, B**).

### Transmembrane stem cell factor-based treatments cause slower c-Kit internalization

SCF interacts with its cognate receptor c-Kit leading to dimerization, signaling and internalization through either a caveolin or clathrin-mediated pathway^38^. We measured the surface c-Kit internalization by flow cytometry and found that tmSCF-based treatments had slower uptake in mast cells (MC9 cells), bone marrow mononuclear cells (BMMNCs) and endothelial cells in comparison to soluble SCF (**Fig. 2C-E**). In endothelial cells, the internalization of c-Kit was the fastest for the tmSCFPLs in comparison to the other tmSCF formulations but still significantly slower than soluble SCF (**Fig. 2E**). Soluble SCF can be internalized by mast cells using a caveolin- or clathrin-mediated pathway^39^. The caveolin-mediated pathway has slower uptake kinetics and leads to reduced proliferation and migration of mast cell^39^. The clathrin-mediated pathway has faster uptake kinetics and leads to increased proliferation and migration of mast cells^39^. To examine the uptake mechanism of the tmSCF formulation, we treated cells with SCF and the tmSCF-based treatments and then performed immunostaining for c-Kit and clathrin or caveolin followed by an analysis for co-localization between the markers. We found that mast cells preferentially use clathrin-mediated pathways to internalize SCF and caveolin-mediated pathways to internalize tmSCFNDs (**Fig. 2F**). Mast cells use both clathrin- and caveolin-mediated pathway to uptake tmSCF or tmSCFPLs. These results correspond well with the protein uptake kinetics results, showing faster uptake of the SCF therapy (a hallmark of clathrin-mediated uptake) and slower uptake of the tmSCF-based therapies (a hallmark of caveolin-mediated uptake; **Fig. 2C**)^39^. Next, we conducted the same experiment on bone marrow derived endothelial progenitor cells (EPCs) and found that they preferentially use clathrin-mediated pathways to uptake SCF while they use a caveolin-mediated pathway to uptake tmSCFPLs and tmSCFNDs (**Fig. 2G**). We found that endothelial cells use both clathrin- and caveolin-mediated pathways to uptake SCF, tmSCF, tmSCFPLs and tmSCFNDs (**Fig. 2H**).

### Transmembrane SCF-based therapies activate the c-Kit pathway in endothelial cells and EPCs but not mast cells

In MC9 mast cells, SCF treatment induced significantly more phosphorylation of c-Kit than tmSCF-based treatments (**Fig. 2F**). In EPCs, SCF and tmSCFND treatment induced significantly more phosphorylation of c-Kit in comparison to control, indicating that SCF and tmSCFNDs may have angiogenic activity related to the SCF/c-Kit pathway (**Fig. 2G**). In endothelial cells, all treatments induced significantly more phosphorylation of c-Kit in comparison to control (**Fig. 2H**). These results support that there is lower pleotropic activity from tmSCF treatments on mast cells. The type of formulation used for delivering tmSCF tailors its trophic specificity, with preferential activation and faster internalization of tmSCFPLs by matured endothelial cells and greater activity towards EPCs using tmSCFNDs.

### Transmembrane SCF proteoliposomes induce greater endothelial cell tube formation in comparison to SCF or tmSCF nanodiscs

To measure the angiogenic activity of tmSCF based therapies, we performed a tube formation assay using endothelial cells. While tmSCF protein itself induced limited tube formation, tmSCFPLs had 16 to 18 fold greater activity (**Supplemental Fig. 3A, B**). In contrast, we did not see a significant induction of tube formation with tmSCFND treatment (**Supplemental Fig. 3C, D**). We also tested endothelial cell migration and proliferation but did not see any significant differences for any of our treatments compared to the control (**Supplemental Fig. 4**).

**Figure 3.**
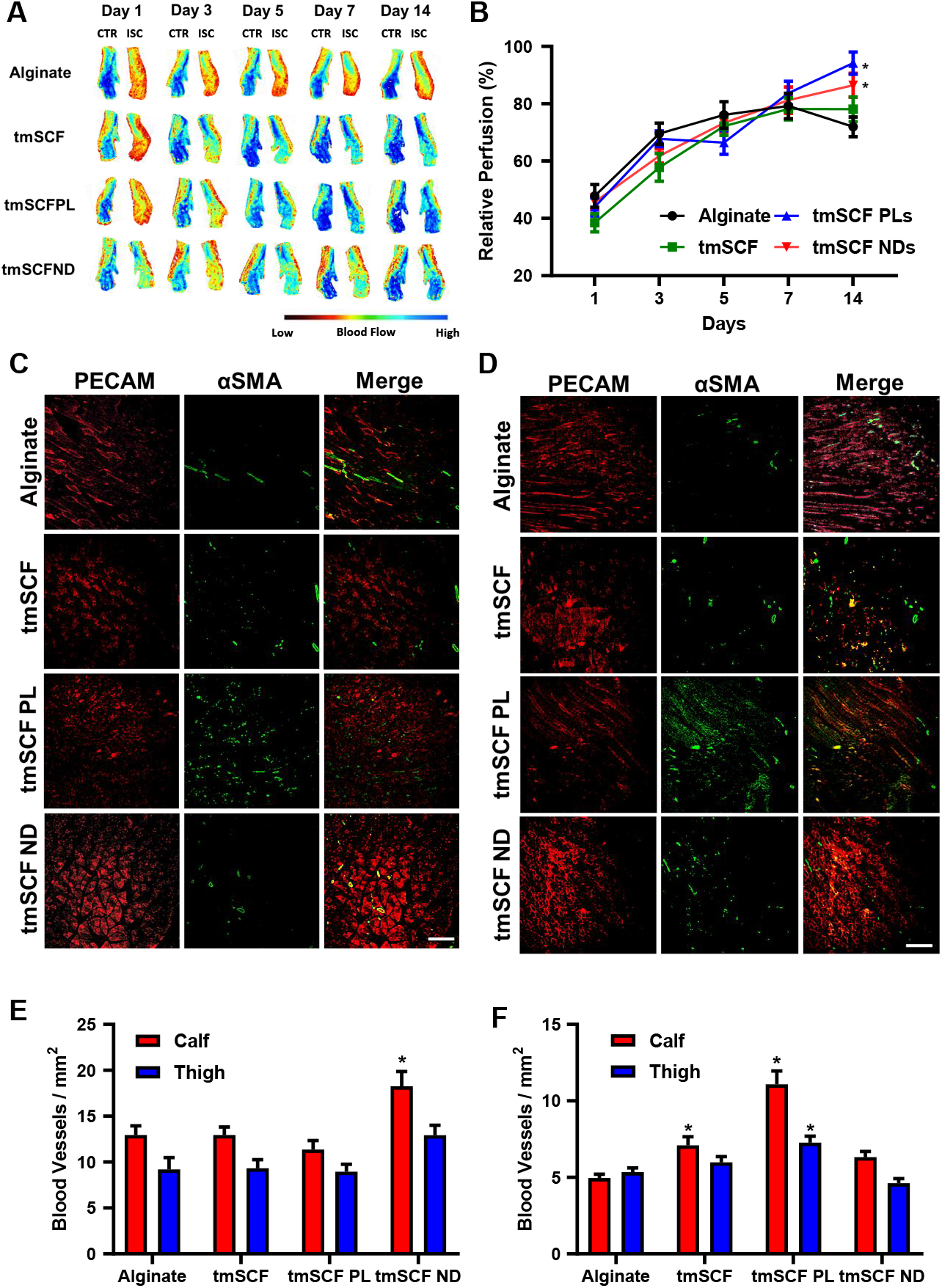
The result of hindlimb ischemia model on wild type mice. (A) Representative mice foot images taken by laser speckle imaging. (B) Relative blood flow recovery after hind limb ischemia surgery on WT mice (n =12-13). (C) Representative immunostaining images of PECAM (red) and αSMA (green) on WT mice calf muscle. Scale bar is 300 μm. (D) Representative immunostaining images of PECAM (red) and αSMA (green) on WT mice thigh muscle. Scale bar is 300 μm. *p<0.05 versus alginate. (E) The number of small blood vessels in the calf and thigh muscle counted from PECAM immunostaining images (n = 4-7). (F) The number of large blood vessels counted from αSMA immunostaining images (n = 4-7).

**Figure 4.**
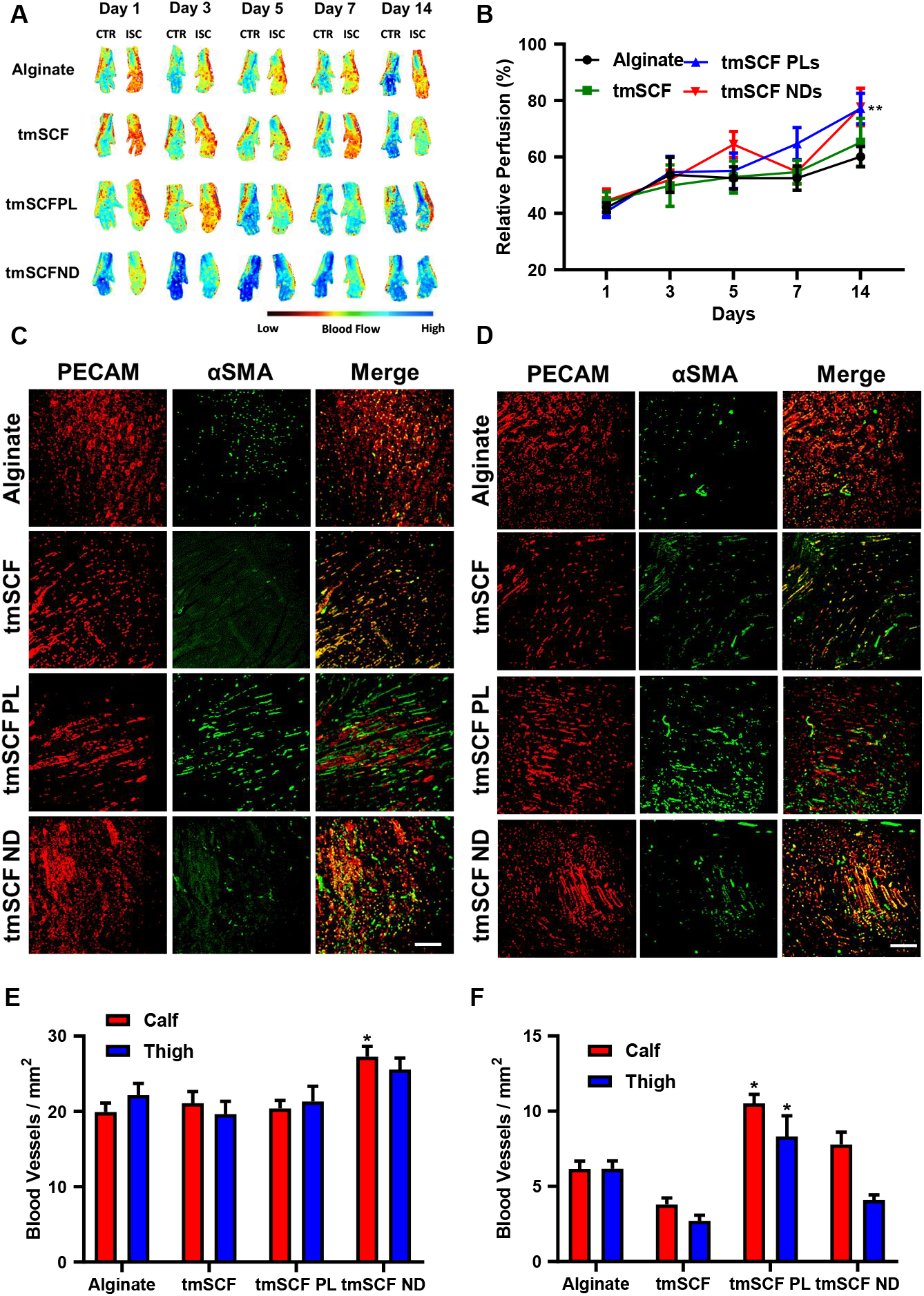
The result of hindlimb ischemia model on ob/ob mice. (A) Representative mice foot images taken by laser speckle imaging. (B) Relative blood flow recovery after hind limb ischemia surgery on ob/ob mice (n = 11-12). (C) Representative immunostaining images of PECAM (red) and αSMA (green) on ob/ob mice calf muscle. Scale bar is 300 μm. (D) Representative immunostaining images of PECAM (red) and αSMA (green) on ob/ob mice thigh muscle. Scale bar is 300 μm. *p<0.05 versus alginate. (E) The number of small blood vessels in the calf and thigh muscle counted from PECAM immunostaining images (n = 4-10). (F) The number of large blood vessels counted from αSMA immunostaining images (n = 4-10).

### Transmembrane stem cell factor proteoliposomes and nanodiscs enhance revascularization in wild type mice

To deliver the treatments *in vivo*, we delivered the treatments from an injectable alginate gel. In vitro measurement of release kinetics demonstrated that the release of tmSCF slightly was faster than for tmSCFPLs or tmSCFNDs (**Supplemental Fig. 5**). The gel was completely degraded after seven day. To examine whether tmSCF-based treatments could enhance revascularization, we ligated the femoral artery of male mice and implanted treatments encapsulated in alginate beads at the ischemic site. We found that there was significantly higher relative blood flow recovery in tmSCFPL/ND groups in comparison to the alginate control group (**Fig. 3A, B**). Immunohistochemical staining for PECAM-1 and α-smooth muscle actin (α-SMA) revealed that there was a significant increase in small blood vessels formed at Day 14 in the tmSCFND treatment group while there was increased mature blood vessels in mice treated with tmSCFPLs (**Fig. 3C-F**).

**Figure 5.**
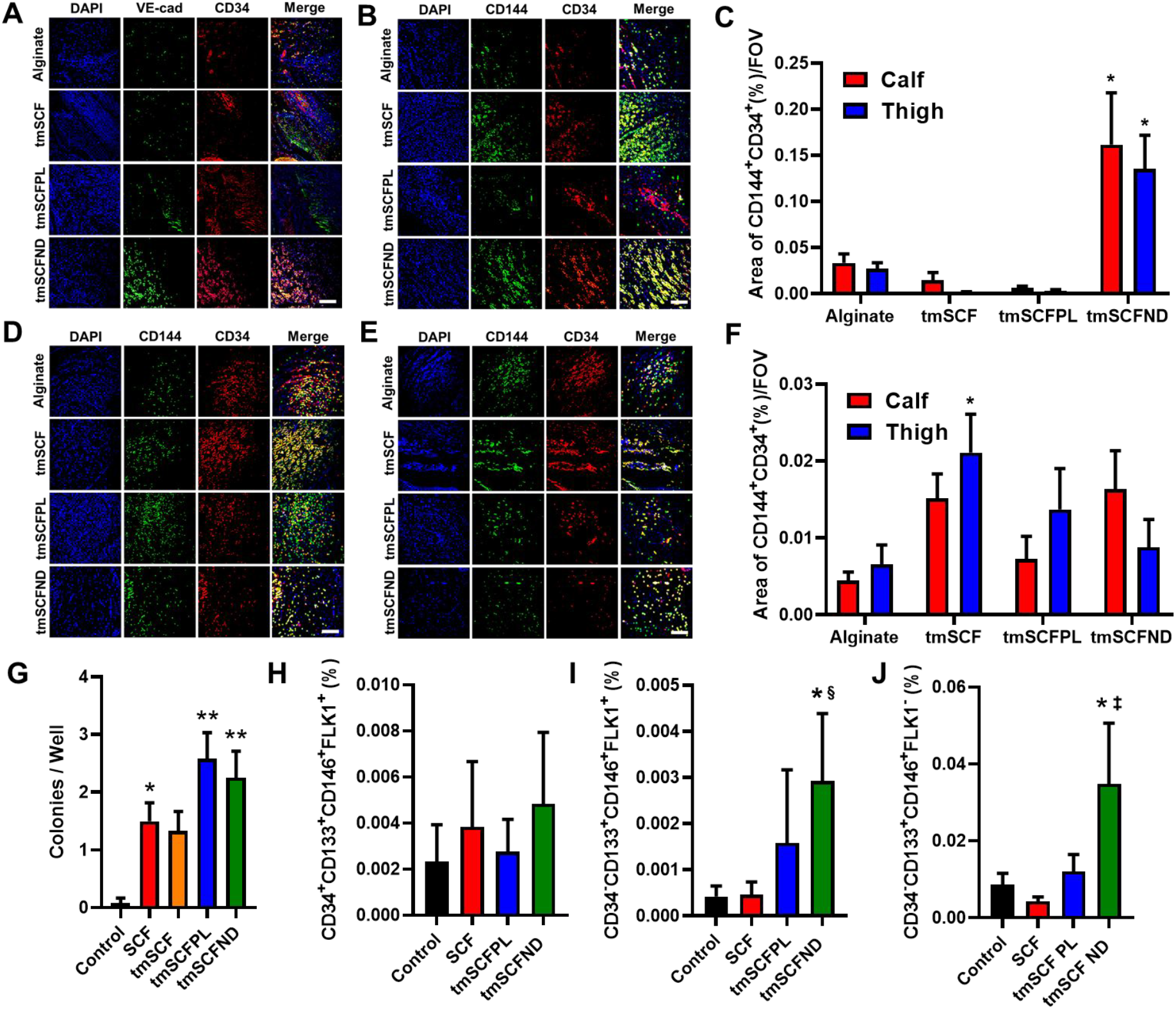
EPCs are recruited to peripheral blood and ischemic site. (A and B) Representative immunostaining images of CD34 (red) and VE-cadherin (green) on WT mice calf and thigh muscle, respectively. Scale bar is 300 μm. (C) CD34 and VE-cadherin double positive areas were quantified in WT mice (n = 3 for thigh, n = 3-4for calf). **p<0.01 versus control (D and E) Representative immunostaining images of CD34 (red) and VE-cadherin (green) on ob/ob mice calf and thigh muscle, respectively. Scale bar is 300 μm. (F) CD34 and VE-cadherin double positive areas were quantified in ob/ob mice (n = 4-10). **p<0.01 versus control. (G) Average large EPC colony number per well (n = 11-12). (H) Frequency of CD34^+^CD133^+^CD146^+^FLK1^+^ cells in peripheral blood after subcutaneous injection (n = 11-12). (I) Frequency of CD34^−^ CD133^+^CD146^+^FLK1^+^ cells in peripheral blood after subcutaneous injection (n = 10-13). (J) Frequency of CD34^−^CD133^+^CD146^+^FLK1^−^ cells in peripheral blood after subcutaneous injection (n = 10-13). *, ^†^, ^‡^ and ^§^ indicate significant difference over alginate control, tmSCF, tmSCFPL, and SCF, respectively (p < 0.05; Kruskal–Wallis one-way analysis of variance).

### Diabetic patients have reduced endothelial stem cell factor expression

Our group and others have identified mechanisms of therapeutic resistance to growth factors^36,40–44^, including the loss of cell surface proteoglycans^36,40–44^ and expression of angiogenesis inhibiting growth factors^45^. Diabetic patients in particular have reduced recruitment of stem cells to ischemic regions, circulating endothelial progenitor cells and endothelial cell function^46,47^. We examined the expression of SCF/tmSCF in skin samples from diabetic and non-diabetic patients to assess differences. We found a marked reduction of SCF expression in endothelial cells in the skin of type 2 diabetic (T2D) patients (**Supplemental Fig. 6A, B**). The expression of c-Kit in endothelial cells was higher in patients with T2D in comparison to non-diabetic patients (**Supplemental Fig. 6C, D**). Conversely, SCF levels were similar between the groups in skin fibroblasts (**Supplemental Fig. 6E, F**). In addition, we found an increase in c-Kit in mast cells in the skin from diabetic patients (**Supplemental Fig. 6G, H**). These results support that there is a significant alteration in SCF and c-Kit in the skin of diabetic patients that may contribute to altered recovery of ischemia.

**Figure 6.**
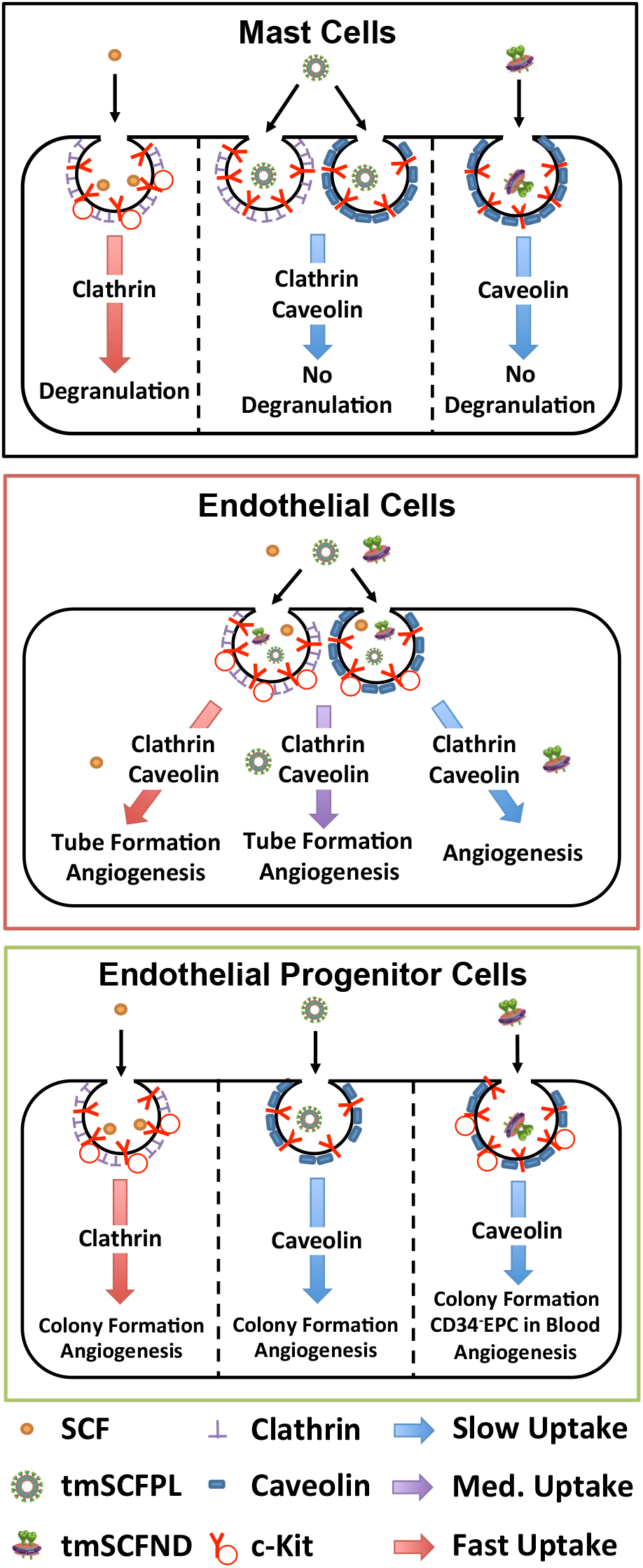
Summary of the experimental findings in the studies. Mast cells use primarily a clathrin-mediated pathway to internalize SCF, leading to mast cell activation and anaphylaxis. In contrast, mast cells use predominantly clathrin and caveolin-mediated pathway to uptake tmSCF-based treatments and these treatments do not cause mast cell activation. Endothelial cells use both of clathrin- and caveolin-mediated pathway to uptake SCF, inducing angiogenesis in endothelial cells. Endothelial cells use both of clathrin- and caveolin-mediated pathway to internalize tmSCFPLs with medium uptake speed, triggering tube formation of endothelial cells and therapeutic angiogenesis. However, tmSCFNDs are internalized through clathrin/caveolin-mediated pathways with slow kinetics and do not induce an angiogenic response from mature endothelial cells. Endothelial progenitor cells (EPCs) use clathrin-mediated pathway to uptake SCF, triggering colony formation of EPCs and bone marrow cell mobilization. For tmSCF-based treatments, EPCs use a caveolin-mediated pathway for internalization leading to colony formation and angiogenesis. Treatment with tmSCFNDs further induced the mobilization of CD34^−^CD133^+^ EPCs to the peripheral blood.

### Transmembrane stem cell factor proteoliposomes and nanodiscs enhance revascularization in ob/ob mice

To examine the effect of the treatments under diabetic conditions, we repeated the hind limb ischemia model in female ob/ob mice. After 14 days, there was significantly higher blood flow recovery in tmSCFPLs and tmSCFNDs treated groups in comparison to the alginate control treated mice (**Fig. 4A, 4B**). Immunohistochemical staining on ob/ob mice to detect endothelial cells (PECAM-1) and smooth muscle cells (α-SMA) showed a increased small blood vessels under tmSCFNDs treatment while there was increased numbers of mature blood vessels under tmSCFPLs treated group (**Fig. 4C-4F**). Treatment with tmSCF alone did not enhance the mature blood vessel formations in ob/ob mice in comparison to WT mice, indicating the better treatment potential with nanocarriers. Overall, tmSCF treatment with nanocarriers improved blood flow recoveries in both WT and ob/ob mice in the hind limb ischemia model.

### Transmembrane stem cell factor nanodiscs induce CD34−CD133+ endothelial progenitor cells mobilization to the peripheral blood

Our trafficking studies supported that the proteoliposomal formulation of tmSCF was preferentially taken up by endothelial cells in comparison to free tmSCF or tmSCFNDs. Consistent with these findings, treatment with tmSCFPLs induced significantly higher tube formation in endothelial cells in comparison to cells with control treatment while tmSCFNDs did not (**Supplemental Fig. 3**). We next treated bone marrow cells for 30 minutes and found that tmSCFPLs and tmSCNDs increased the population of CD34^−^CD133^+^CD146^+^ cells in comparison to the control or tmSCF treated cells (**Supplemental Fig. 7A**). A recent study identified that CD34^−^CD133^+^ EPCs were the most potent EPC phenotype for treating ischemia in the recent study^48^, suggesting that tmSCFPLs and tmSCFNDs specifically induced this potent EPC phenotype. We immunostained for CD34^+^/CD144^+^ in the muscles of WT mice with hind limb ischemia and found increased EPCs in the ischemic muscles of tmSCFNDs treated mice, indicating the recruitment of EPCs to the ischemic region (**Fig. 5A-C**). We also immunostained for EPCs in the muscles of ob/ob mice from the hind limb ischemia study and found increased CD34^+^/CD144^+^ in the ischemic muscles of tmSCF treated mice (**Fig. 5D-F**). We also conducted an EPC colony formation assay on BMMNCs and found that both tmSCFPLs and tmSCFNDs induced a greater number of large EPC colonies in comparison to the control (**Fig. 5G**). Together, these results suggest that nanocarrier formulations of tmSCF are more active in inducing bone marrow cell mobilization and EPC recruitment in comparison to the free tmSCF.

Following injury or development of ischemia, EPCs mobilize and home to the region of injury^49^. Soluble SCF is known to participate in both the mobilization and homing of EPCs in concert with other cytokines^50^. To test if the tmSCF-based treatments induced EPC mobilization, we performed subcutaneous injections of the treatments in mice for four days consecutively and then performed flow cytometry on the peripheral blood to identify EPCs. While we did not observe significant changes in the conventional EPC population of CD34^+^CD133^+^CD146^+^FLK1^+^ cells (**Fig. 5H**), we found that there were increases in CD34^−^CD133^+^CD146^+^FLK1^+^ and CD34^−^CD133^+^CD146^+^FLK1^−^ cells in the tmSCFND treated group (**Fig. 5I, J**). This result indicates that tmSCFNDs induce mobilization of the potent CD34^−^ subtype of EPCs to peripheral blood as described in previous work^48^. We also examined the EPC population in bone marrow after four consecutive days of subcutaneous injection of our treatments and found no significant alterations in the CD34^−^CD133^+^CD146^+^FLK1^+^ or CD34^+^CD133^+^CD146^+^FLK1^+^ populations in the bone marrow following the treatments **(Supplemental Fig. 7B, C)**.

## Discussion

Overall, our study demonstrated that tmSCF is an effective therapeutic protein for enhancing revascularization in peripheral ischemia without the accompanying activation of mast cells. Membrane proteins have been traditionally viewed as targets for drug development and are often difficult to isolate and maintain in solution in the absence of the cell membrane. Our work demonstrates that the transmembrane form of SCF may provide advantages over the soluble SCF, with reduced effects on mast cells while maintaining activity to induce bone marrow cell mobilization and angiogenesis. The treatment of ischemia in diabetes has been a recurrent clinical problem, especially among diabetic patients receiving bypass surgery or percutaneous interventions^51,52^. Our work has demonstrated that there are alterations in SCF and c-Kit expression in diabetic patients and that tmSCF-based therapeutics are effective in treating ischemia in diabetic mice. Additionally, our work supported that the nature of the lipid nanocarrier used to encapsulate tmSCF can significantly alter its interactions with different cell types, leading to alterations in the regeneration process during revascularization in ischemia.

A major difference in the tmSCF-based treatments in comparison to soluble SCF was the slower uptake kinetics and cellular trophism of tmSCF for endothelial cells or EPCs depending on the nanocarrier used for delivery. A summary of the findings of our trafficking studies is shown in **Fig. 6**. Mast cells use predominantly a clathrin-mediated pathway to internalize SCF, clathrin and caveolin-mediated pathways for tmSCFPLs, and a caveolin-mediated pathway for tmSCFNDs (**Fig. 6**). The clathrin-mediated pathway has more rapid internalization and causes increased activation of mast cells^39^. Our studies suggest that mast cell activation does not occur for tmSCF-based treatments because of the slower uptake, greater utilization of the caveolin internalization pathway and weaker activation of the c-Kit receptor. A recent study used protein engineering techniques to create a modified version of SCF that led to partial agonism of the c-Kit receptor, allowing for therapeutic activity towards hematopoietic stem cells without mast cell activation^53^. These findings are consistent with our studies that show weaker activation of c-Kit with reduced mast cell activation and therapeutic activities.

Our studies also revealed that the type of nanocarrier used to deliver tmSCF elicited a differential response based on cell type. Endothelial cells were most responsive to tmSCF proteoliposomes, showing increased tube formation, angiogenesis and mature vessel formation (**Fig. 6**). In contrast, EPCs were more responsive to tmSCF nanodiscs and induced greater colony formation and mobilization of CD34^−^CD133^+^ EPCs to the peripheral blood. Our studies suggest that endothelial cells use a combination of clathrin and caveolin-mediated pathways to internalize tmSCFPLs and tmSCFNDs while EPCs use only a clathrin-mediated pathway to internalize these treatments. Notably, the size of proteoliposomes used here was chosen as it was the optimal size for delivering other therapeutic membrane proteins to induce angiogenesis and endothelial cell activation in our previous studies^36,40,41,44^. One potential explanation for the differences in uptake mechanism between the treatments may be that the size of the therapeutic compounds. A previous study demonstrated that cells use predominantly clathrin-mediated pathways to internalize particles <200 nm in size while they use almost exclusively caveolin-mediated pathways to take up particles >500 nm^54^. In our study, the size of the tmSCFNDs was around 200 nm and the size of the tmSCFPLs was approximately 400 nm. Thus, our findings are consistent with this previous work and suggest that formulating membrane protein therapeutics spanning these ranges is an effect means for controlling the therapeutic response if there are clathrin and caveolin-based difference in the cellular response to the protein. In addition, these findings provide a potential explanation for the cell type specific response as mast cells, endothelial cells and EPCs could respond differentially based on their differences in the utilization of these pathways for uptake of the c-Kit receptor.

In summary, this study demonstrates the first use of tmSCF as a therapeutic protein for the treatment of peripheral ischemia. Our work supports that there is minimal mast cell activation by tmSCF-based therapies and that the activity can be targeted to specific cell types and internalization pathways through the use of different nanocarriers. While both nanodisc and proteoliposome-delivered tmSCF are beneficial in treating ischemia, our work supports that the proteoliposomal tmSCF may act through the stimulation of angiogenesis in endothelial cells and proliferation of bone marrow cells. Conversely, nanodisc delivered tmSCF acts primarily through the mobilization and recruitment of bone marrow cells and stimulation of a CD34^−^ EPC population. Overall, our work suggests some of the benefits of SCF treatment can be recapitulated using tmSCF therapies and these appear to have improved safety as well as tailorable activity based on the lipid nanocarrier used.

## Materials and Methods

### Preparation of tmSCF Proteoliposomes

For the production of recombinant tmSCF, HEK-293Ta cells were transduced with lentiviral vectors with constitutive expression of 6X His tagged tmSCF. Viruses were produced in HEK-293Ta cells using human lentiviral packaging system according to the manufacturer’s instructions (Genecopoeia). Puromycin was used to select only for transduced cells. The cells were lysed in a buffer containing 20mM Tris (pH 8.0), 150mM NaCl, 1% Triton X-100, 2mM sodium orthovanadate, 2mM PMSF, 50mM NaF, and protease inhibitors (Roche) at room temperature. Transmembrane stem cell factor (tmSCF) was then isolated using cobalt chelating column (Chelating column; GE Healthcare), and the buffer was exchanged in 1X PBS. After concentrating the protein solution with Centriprep concentrators (Millipore), the final working concentration of protein was found to be 100 μg/ml using a BCA Protein Assay kit (Thermo Scientific). Purity of the protein was confirmed by SDS-PAGE followed silver staining of the gel. To prepare liposomes, stock solutions (10 mg mL^−1^) of 1,2-dioleoyl-sn-glycero-3-phosphocholine, 1,2-dioleoyl-sn-glycero-3-phosphoethanolamine, cholesterol, and sphingomyelin were prepared in chloroform. The lipids were mixed in the volumetric ratio 2:1:1:1 in a round bottom flask and the chloroform was removed on rotatory evaporator attached to a vacuum pump. Liposomes were resuspended in 4-(2-hydroxyethyl)-1-piperazineethanesulfonic acid buffer by vortexing, sonication, and freeze thawing. Extruding device (Avanti Polar Lipids) with polycarbonate membranes (400 nm) was used to homogenize the liposome population. A mild detergent, n-octyl-β-D-glucopyranoside (1% w/v) was added to both the liposome and the recombinant tmSCF. The protein and liposomes were then combined and the detergent was removed by timed serial dilution (every 30 min, 10% dilution up to 2h), dialysis, and treatment with Biobeads (Biorad). The final tmSCF concentration in tmSCFPLs and tmSCFNDs were determined 55 μg/ml and 77 μg/ml, respectively.

### Preparation of tmSCF Nanodiscs

As a lipid source, 1-palmitoyl-2-oleoyl-sn-glycero-3-phosphocholine (POPC) was used in this study. POPC was stored in chloroform, so we first removed chloroform by rotary evaporator. After that, POPC was resuspended in sodium cholate (100 mM). MSP protein was then added to phospholipid solution, and the detergent concentration was adjusted between 14-40 mM. This construct was incubated for 15 minutes at 4°C. To solubilize the membrane protein, tmSCF was incubated in the n-octyl-β-D-glucopyranoside (1% w/v) for 15 minutes at 4°C. After 15 minutes incubation of lipid construct and tmSCF protein, these were combined and incubated for 1 hour at 4°C. Final detergent concentration was adjusted to 20 mM with sodium cholate. Finally, detergent was removed by dialysis and biobeads.

### Liposome characterization

The size and dispersion of the proteoliposomes were characterized by dynamic light scattering (Malvern Zetasizer Nano ZS). Calibration was performed using 54 nm polystyrene particles. For TEM imaging of proteoliposomes, carbon support grids (300 mesh Cu; EM Sciences) were treated with glow discharge at 50 mA for two minutes (Emitech K100X; Quorum Technologies). The samples were then applied to the grids and the excess liquid removed with a filter paper. 2% uranyl acetate solution was used to stain glids, and images were taken using an FEI Tecnai Transmission Electron Microscope (TEM). For cryo-electron microscopy imaging, the liposomes were plunge-frozen in liquid ethane on carbon holey film grids (R2X2 Quantifoil; Micro Tools GmbH, Jena, Germany). The grids were transferred to a cryo-specimen holder (Gatan 626) under liquid nitrogen and put in a microscope (JEOL 2100 LaB6, 200 keV), Grids were maintained at close to liquid nitrogen temperature during EM session (−172 to −180°C).

### tmSCF protein release kinetics from alginate beads

Purified tmSCF protein was first labeled with Alexa Fluor^™^ 488 Microscale Protein Labeling Kit (Thermo Scientific), then tmSCFPLs and tmSCFNDs were fabricated using this labeled protein. Equal volume of treatments and 4% sodium alginate solution were mixed, then the solution was extruded from 30G needle into a 1.1% calcium chloride solution to crosslink for 1 hour. Protein amount was calibrated based on the protein fluorescent signal. Fabricated alginate beads were incubated in the 5 ml of 1% BSAPBS in plastic scintillation vial and incubated at 37°C. 100 μl of released protein was collected each time point and fluorescent intensity was measured by plate reader. The same amount of 1% BSAPBS was replaced each time point to compensate.

### Tube formation assay

Cells were stained with cell tracker green, cultured and starved 24h prior to the experiment. Growth factor reduced Matrigel (Corning) were coated on the 96 well plate, and endothelial cells were seeded on top of it with seeding density of 4×10^4^/well. Treatments were then added and cells incubated for 6 or 10 hours. Images were taken using high throughput imaging system (Cytation 5 Cell Imaging Multi-Mode Reader; Biotek) at each time point and the number of loop was quantified using Photoshop (Adobe).

### Cell proliferation assay

Endothelial cells were passaged into a 96-well plate and cultured in low serum media with 2% FBS for 24h. Treatments were then added to cells. After 24h, BrdU was added to the cells and proliferation was assessed 12h later using a colorimetric BrdU assay (Cell Signaling, Inc.).

### Cell migration assay

Endothelial cells were stained with cell tracker green, and passaged to confluence in 96-well migration assay kit (Platypus). Treatments were added for 24h before assay starts. A cell stopper was then removed to allow cells to migrate. Migration area and green fluorescent intensity were measure after 16 and 24 hours.

### Hind limb ischemia model

Prior to surgery, alginate beads containing the treatments were fabricated by mixing equal volume of 4% sodium alginate solution and treatments. Beads were created through extruding the alginate solution through a 30G needle into a 1.1% calcium chloride solution to crosslink for 1 hour. Wild-type mice and ob/ob mice (n = 12-13 for WT, n = 11-12 for ob/ob; Jackson Labs) were used in the studies. To perform the hind limb ischemia studies, mice were anesthetized with 2-3% isoflurane gas and a longitudinal incision was made in the inguinal region of the right thigh. The femoral artery was separated from the femoral vein and nerve, and then double ligated with 6-0 silk sutures at two locations and the artery severed at each ligation. Treatments were then implanted in the pocket created by the surgery and the wound was closed with resorbable sutures (4-0 Vicryl; Ethicon, Inc.).

### Immunohistochemistry on human skin

Human skin samples were obtained from the Glasgow Caledonian University Skin Research Tissue Bank, Glasgow UK. The tissue bank has NHS research ethics approval to supply human skin for research (REC REF: 16/ES/0069). All methods were carried out in accordance with relevant guidelines and regulations. All experimental protocols were approved by the NHS East of Scotland Research Ethics Service. Informed consent was obtained from all subjects (no patients were under 18 years of age).

Samples were formalin fixed and embedded in paraffin following standard procedures prior to sectioning. The sections were then deparaffinized in xylene and treated for 3 hours with antigen retrieval solution (Dako) at 80°C. The sections were cooled to room temperature and blocked with 20% fetal bovine serum for 45 minutes and then immunostained overnight with 1:100 dilution of primary antibody to PECAM-1 (Cell Signaling), SCF (Abcam), FSP1 (Abcam), c-Kit (Abcam), and mast cell tryptase (Abcam). Secondary staining was performed with secondary antibodies conjugates with Alexa Fluor 488 or 594 dye (Thermo Scientific). Following staining, the samples were imaged using confocal microscopy. For quantification, double-positive areas were quantified using Photoshop and ImageJ.

### Immunohistochemistry on mice

The thigh and calf muscles were formalin fixed and embedded in paraffin following standard procedures prior to sectioning. The sections were deparaffinized in xylene and treated for 3 hours with antigen retrieval solution (Dako) at 80°C. The sections were cooled to room temperature and blocked with 20% fetal bovine serum for 45 minutes and then immunostained overnight with 1:100 dilution of primary antibody to PECAM-1 (Cell Signaling), αSMA (Abcam), CD34 (Santa Cruz), and VE-cadherin (Abcam). Secondary staining was performed with secondary antibodies conjugates with Alexa Fluor 488 or 594 dye (Thermo Scientific). Following staining, the samples were imaged using confocal microscopy. The number small blood vessels was counted from PECAM-1 positive area, and the number of matured blood vessels was counted from αSMA positive area. For the quantification of the double-positive areas of CD34 and VE-cadherin were quantified using Photoshop and ImageJ.

### EPC colony formation assay

Bone marrow cells (BMCs) were harvested, and Bone marrow mononuclear cells (BMMNCs) were then separated by using Ficoll density centrifugation. Semisolid culture medium was prepared by mixing methylcellulose (MethoCult^™^ SF M3236; Stem Cell Technologies), 15% fetal bovine serum, 50 ng/ml murine vascular endothelial growth factor, 50 ng/ml murine basic fibroblast growth factor, 50 ng/ml murine insulinlike growth factor-1, 20 ng/ml murine interleukin-3, 50 ng/ml murine epidermal growth factor, and 100 ng/ml stem cell factor in IMDM. SCF is replaced with the same concentration of tmSCF, tmSCFPL, and tmSCFND respectively for each treatment group. Both of the BMCs and BMMCs are adjusted to 7×10^5^ cells / ml, then resuspended in the semisolid culture medium. Cell medium is added to 6 well cell culture plate and incubated for 7 days. The number of small and large colony was counted by two investigators who were blinded to the experimental conditions.

### Bone marrow cell induction to CD34− CD133+ CD146+ EPCs

BMCs were harvested freshly for the experiment and cultured in DMEM high glucose media with 10% FBS. 100 ng/ml of SCF, tmSCF, tmSCFPLs, tmSCFNDs were added to the BMCs media, and cells were incubated for 30 minutes. Treatment was stopped by placing the cell plate on ice. Then cells are stained for CD34−Alexa647 (BD Biosciences), CD133-PE (BioLegend) and CD146-Alexa488 (BD Biosciences). CD34−CD133+CD146+ cells are quantified by FlowJo software.

### EPCs induction to peripheral blood

To measure the ability of treatments to induce EPCs to peripheral blood, 240μg/kg of SCF, tmSCF, tmSCFPLs, and tmSCFNDs were injected subcutaneously for consecutive 4 days. At the end of day 4, peripheral blood was collected for EPC marker analysis. As EPCs markers, flk1, CD146, CD34, and CD133 were used for flow cytometry analysis.

### Evans Blue extravasation assay

Male C57BL/6J wild-type mice were anesthetized with 2 % isoflurane and initial ear thickness was measured using Mitsutoyo Micrometer (Uline). 250 mL sterile-filtered 1% Evans blue (Sigma) in PBS was injected into the retro-orbital vein and allowed to recover. After 15 minutes later, mice were anesthetized again and intradermally injected into the ear pinnae with 25 mL of PBS alone (into the left ear pinna) or containing 50 mg/mL SCF, tmSCF, tmSCFPLs, or tmSCFNDs (into the right ear pinna) using a 1 mL syringe equipped with a 30 G × 1/2 needle. Two and three hours after injections, mice were anesthetized again and ear pinnae thickness was measured and photographs were taken. Three hours after injections, mice were euthanized, ear pinnae were harvested. Ear pinnae was incubated in 300 mL dimethyl sulfoxide (DMSO) in a 48-well tissue culture plate for 20 hours on a shaker at room temperature. Evans blue containing supernatant was collected, transferred into 96-well flat-bottom plate and absorbance at 620 nm wavelength was measured using a plate reader.

### C-Kit internalization by flow cytometry

MC9 mast cells, bone marrow cells, and HUVECs were starved overnight with 0.5% serum prior to the experiment. The cells were treated with Brefeldin A (5 μg/ml; Cayman Chemicals, Inc.) for 2 hours to inhibit a repopulation of surface c-Kit. The cells were washed with 37C° HEPEs buffer (10 mM HEPES, 137 mM NaCl, 2.7 mM KCl, 0.4 mM Na2HPO4⋅7H2O, 5.6 mM glucose, 1.8 mM CaCl2⋅2H2O, and 1.3 mM MgSO4⋅7H2O with 0.04% BSA) and resuspended at 5 × 10^5^ cells/ml in HEPES buffer without BFA. The cells were then treated with either of SCF, tmSCF, tmSCFPLs, and tmSCFNDs (100 ng/ml) for 5, 10, 20, and 30 minutes. After the treatment, cells were plunged into ice to stop the c-Kit internalization. For HUVECs, accutase was used to detach cells. The surface c-Kit was stained with PE-Cy^™^7 anti-Mouse CD117 antibody (BD Biosciences) or PE-Cy7 anti-Human CD117 antibody (Biolegend) then surface c-Kit intensity was read by flow cytometry. The surface c-Kit intensity was normalized to initial time point (0 min).

### Clathrin, caveolin and c-Kit colocalization assay

Prior to the experiment, bone marrow cells were harvested from 10 weeks old female C57BL/6J wild-type mice and mononuclear cells are isolated by Ficoll density centrifugation. These bone marrow mononuclear cells are cultured for seven days under endothelial media to induce spindle shaped adherent EPCs. The EPCs, MC9 mast cells, and HUVECs were starved overnight with 0.5% serum prior to the experiment. 100 ng/ml of SCF, tmSCF, tmSCFPLs and tmSCFNDs were then treated for two minutes and immediately plunged into ice to stop the internalization. After washing with PBS, and fixing with 4% PFA, immunostaining with clathrin (Abcam), caveolin (Abcam) and c-Kit(Invitrogen, NOVUS Biological) were conducted. Images of 30 cells are taken by confocal microscopy and we analyzed the colocalization of clathrin and c-Kit and caveolin and c-Kit by coloc 2 in Image J. Pearson’s R value was used for analysis.

### Immunostaining

Prior to the experiment, the cells were starved overnight with 0.5% serum prior to the experiment. The cells were then treated with 100 ng/ml of SCF, tmSCF, tmSCFPLs or tmSCFNDs for five minutes and immediately plunged into ice to stop the phosphorylation. After washing with PBS, and fixing with 4% PFA, immunostained for phospho-c-Kit (R&D) was conducted. Images of 30 cells are taken by confocal microscopy and we analyzed the mean intensity of phospho-c-kit inside of a cell.

### Statistical analysis

All results are shown as mean ± standard error of the mean. Comparisons between only two groups were performed using a two-tailed Student’s t-test. Differences were considered significant at p<0.05. Multiple comparisons between groups were analyzed by two-way ANOVA followed by a Tukey post-hoc test. A two-tailed probability value <0.05 was considered statistically significant.

## Supplemental Figure Legends

**Supplemental Figure 1.**
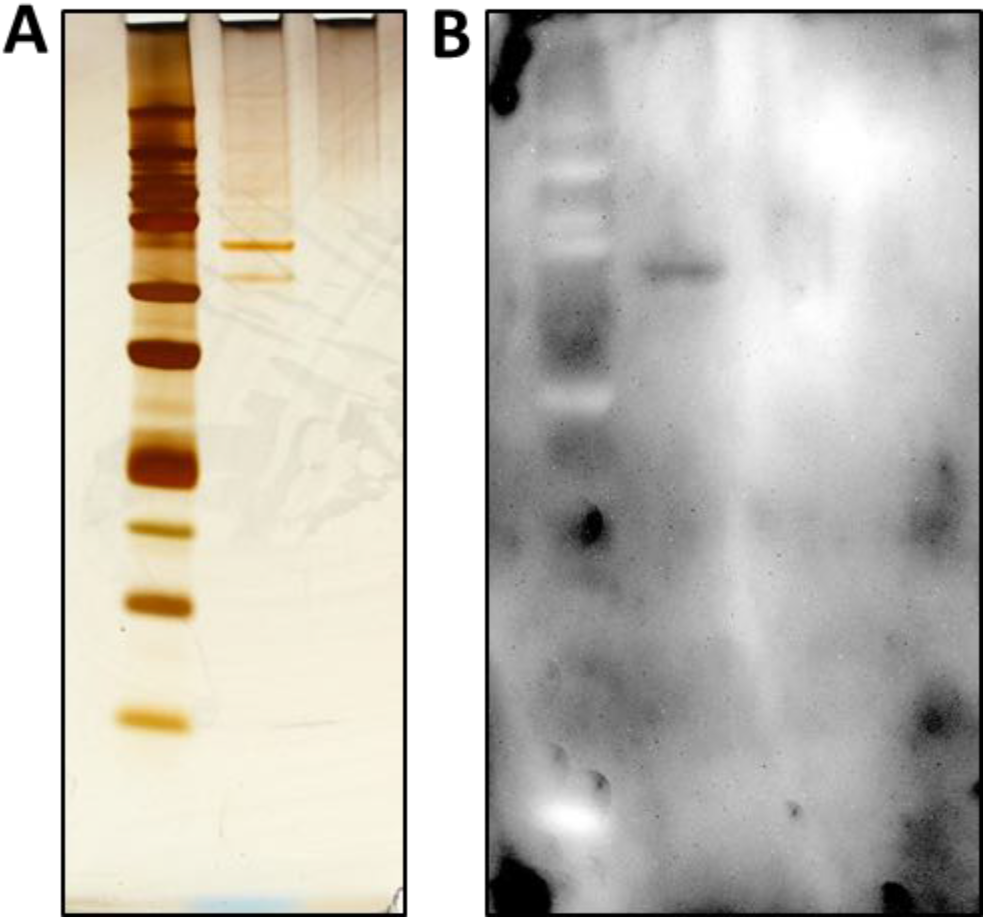
Transmembrane SCF protein purification. (A) Western Blot result of purified tmSCF protein, showing its corresponding band. (B) Silver staining result of purified tmSCF protein.

**Supplemental Figure 2.**
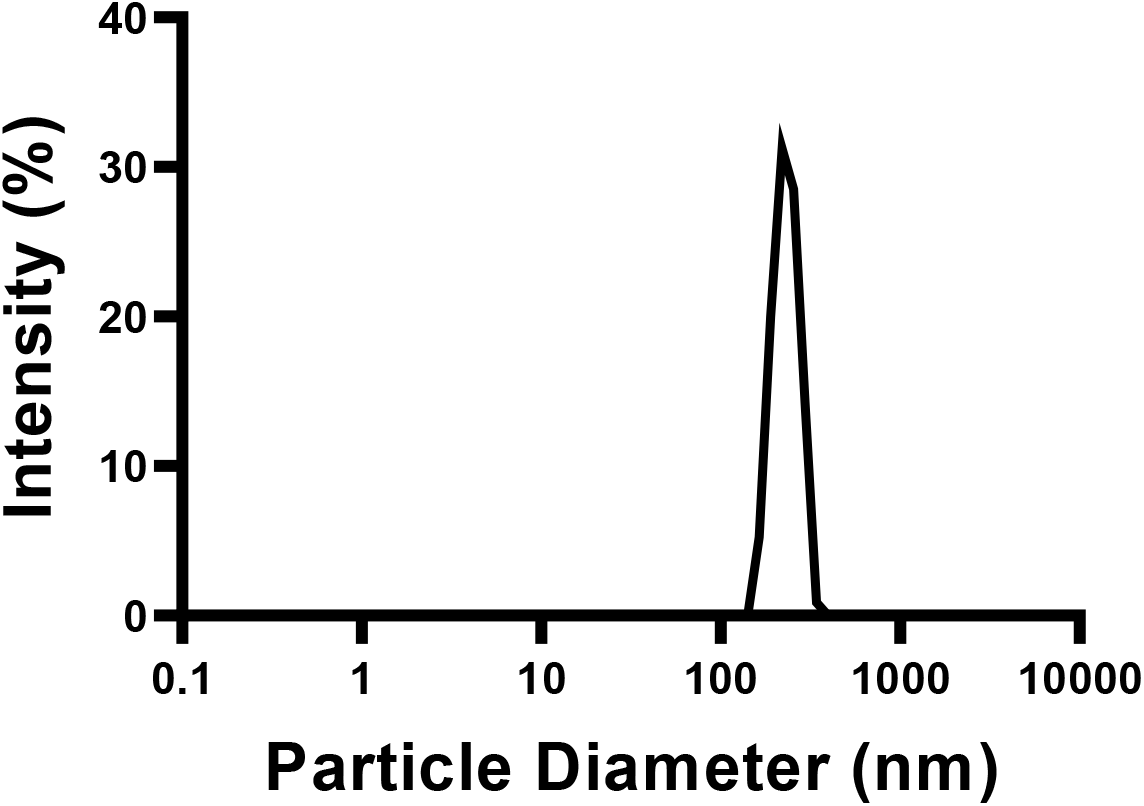
DLS measurement of purified tmSCF protein. Size distribution for tmSCF measured by dynamic light scattering. The average size of tmSCF was approximately 200 nm.

**Supplemental Figure 3.**
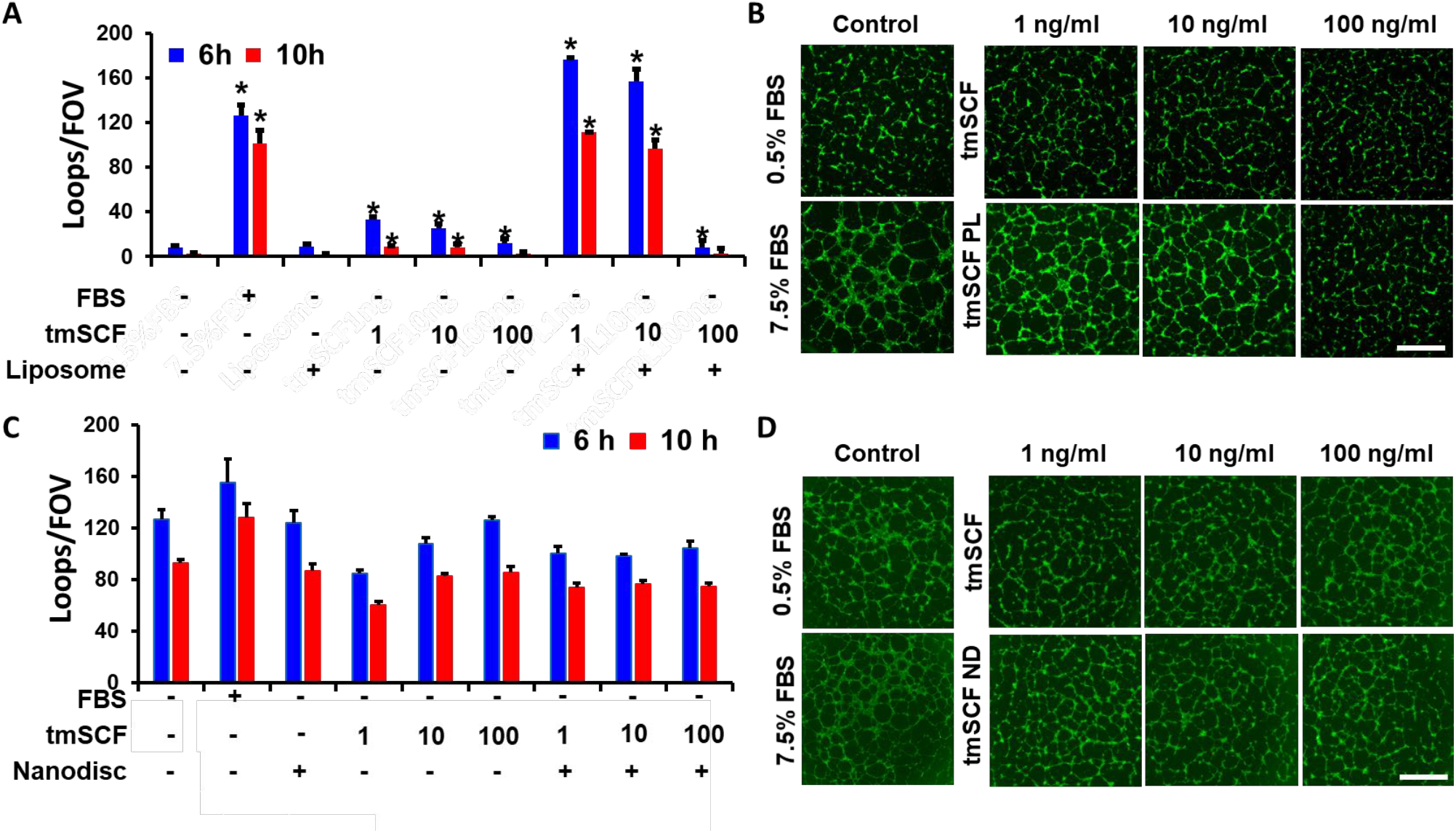
Endothelial cell tube formation assay. (A) HUVECs were starved in MCDB131 (Thermo Scientific) supplemented with 0.5% FBS, L-glutamine and Penicillin-Streptomycin 24 hours prior to the experiment and treated with tmSCF or tmSCFPLs. Loop number was counted after 6 and 10 hours of incubation. Significantly higher number of loops were formed in 1 ng/ml and 10 ng/ml concentration of tmSCFPLs groups compared to negative control. (B) Representative images of HUVECs after 6 hours of tmSCF or tmSCFPLs treatment. Scale bar is 300 μm. (C) HUVECs were treated with tmSCF or tmSCFNDs. No significant difference was confirmed on any of the treatment group. (D) Representative images of HUVECs after 6 hours of tmSCF and tmSCFNDs treatment. Scale bar is 300 μm.

**Supplemental Figure 4.**
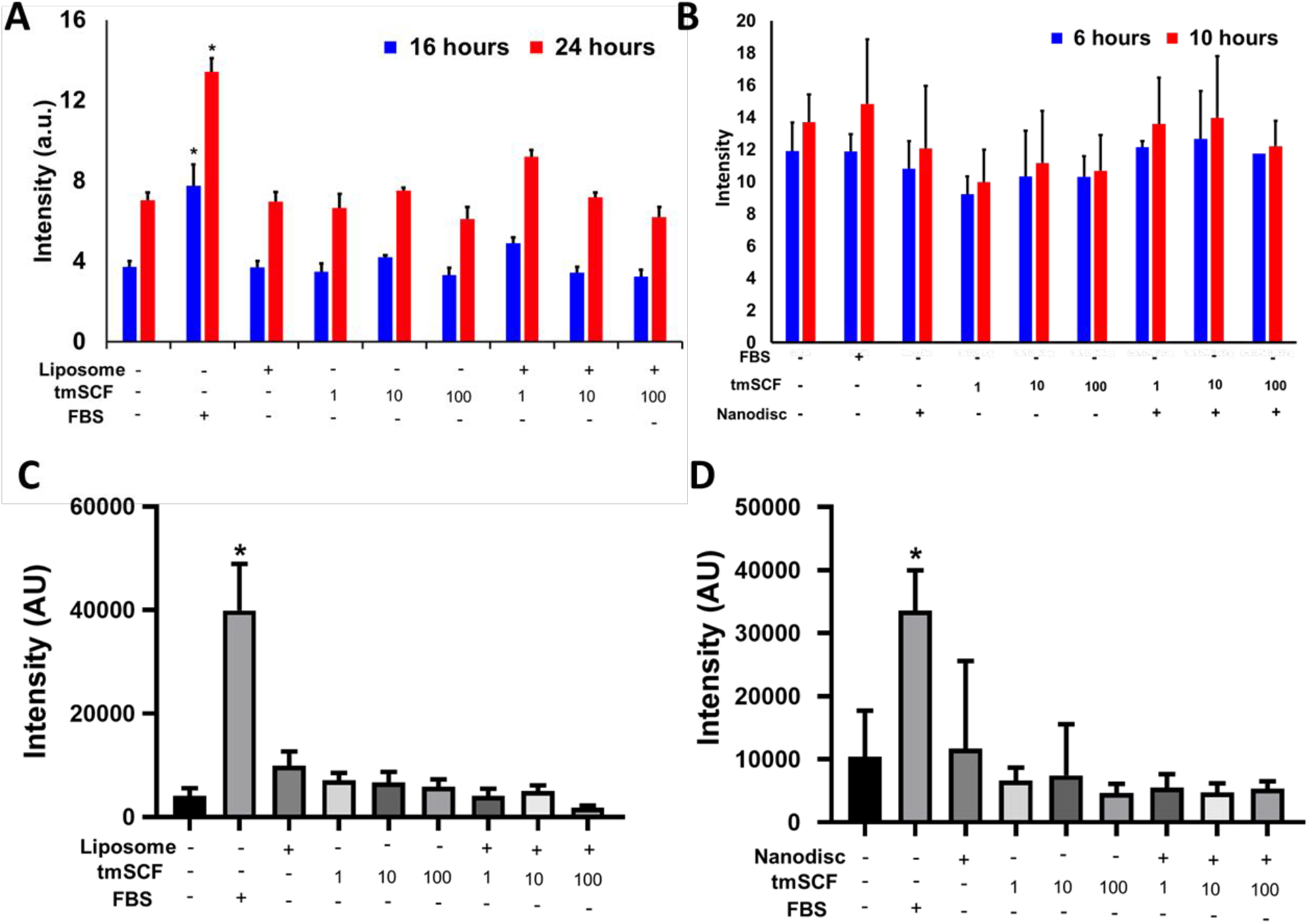
Endothelial cell migration and proliferation assay. (A) Quantification results of migration assay, HUVECs were treated by tmSCFPLs. The number of the cells migrated towards the center were counted by green fluorescent signal. (B) Quantification results of migration assay, HUVECs were treated by tmSCFNDs. The number of the cells migrated towards the center were counted by green fluorescent signal. (C) BrdU intensity was measured after HUVECs were treated by tmSCFPLs. (D) BrdU intensity was measured after HUVECs were treated by tmSCFNDs.

**Supplemental Figure 5.**
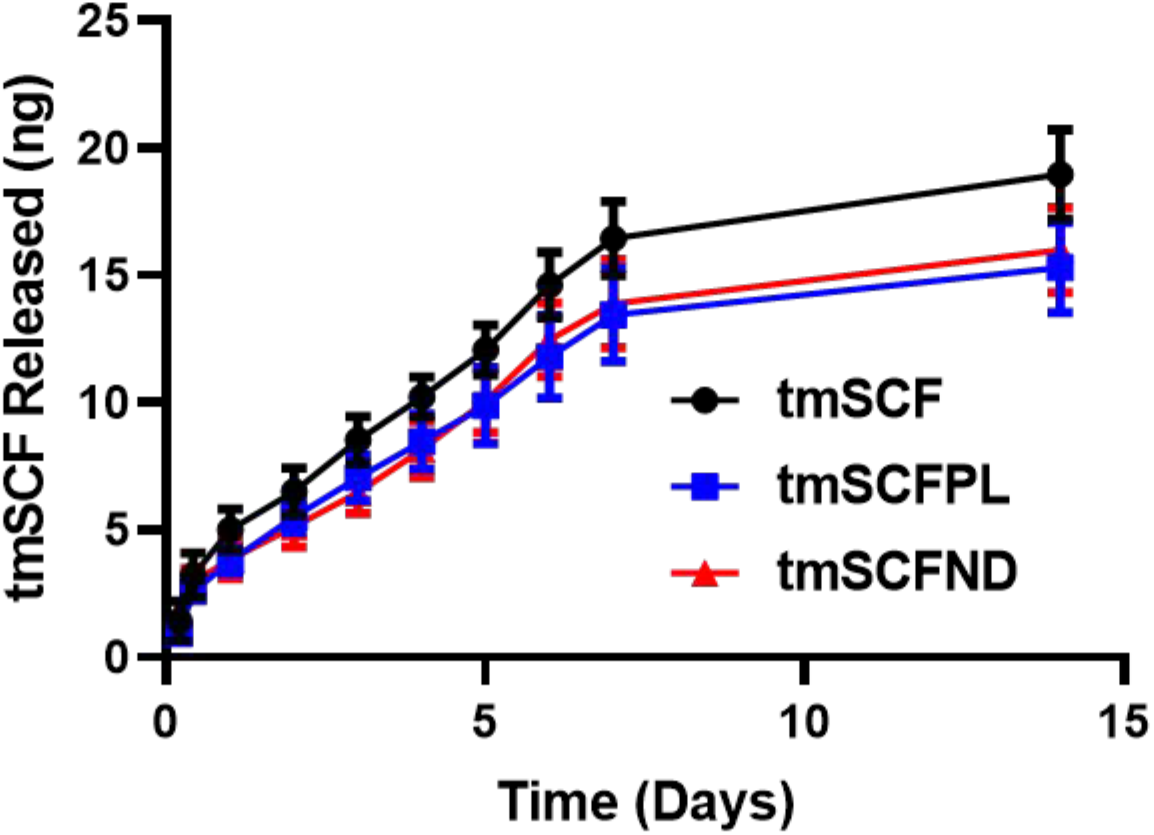
Transmembrane SCF release kinetics from alginate gel showed fast release of tmSCF in comparison to tmSCFPLs or tmSCFNDs. The release kinetics of tmSCF protein, tmSCFPLs, and tmSCFNDs from alginate beads over time.

**Supplemental Figure 6.**
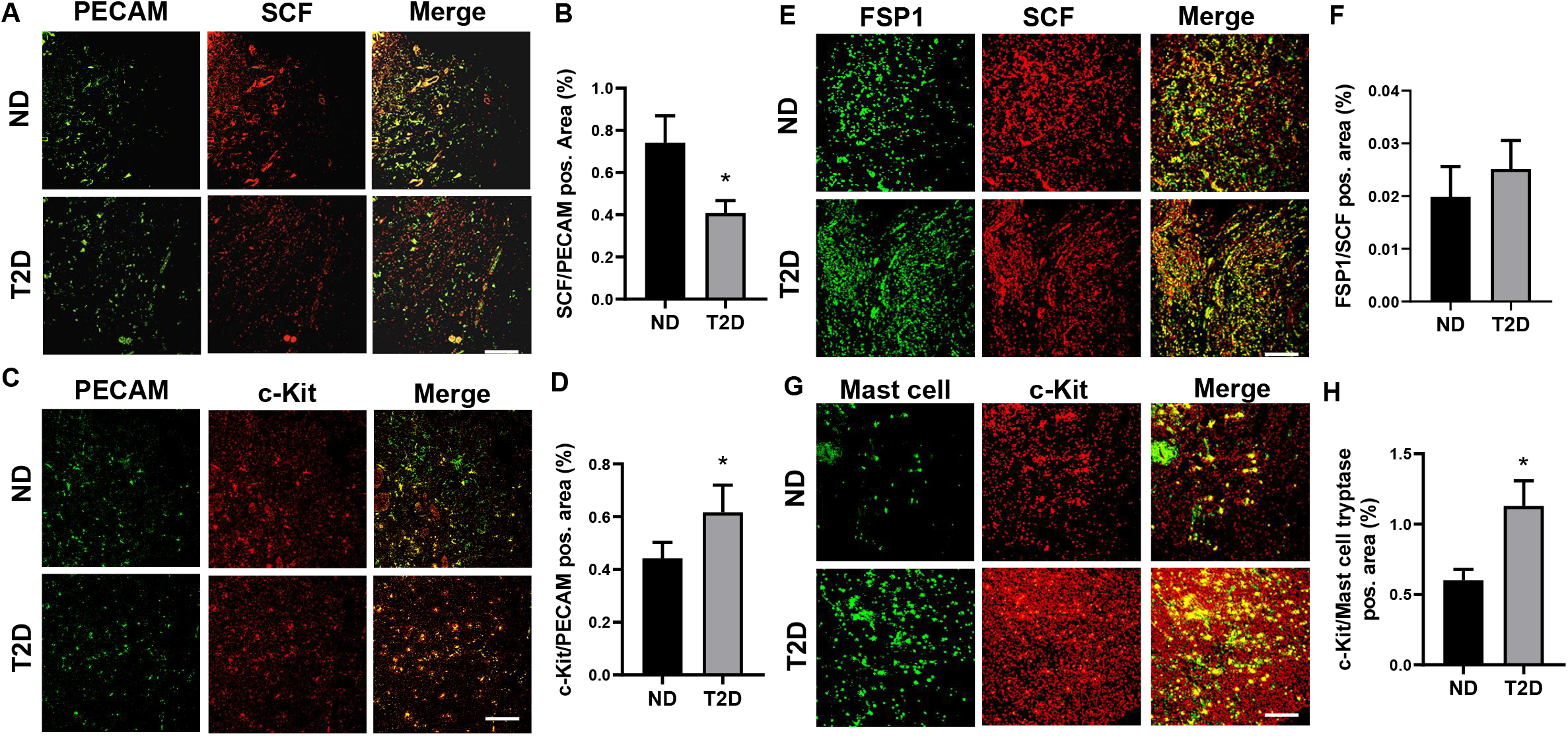
Immunostaining on the skins of patients with diabetes. (A) Representative immunostaining images for PECAM and SCF. Scale bar is 300 μm. (B) Quantification of area of double positive area for PECAM and SCF. *p<0.05 versus nondiabetic group (n = 6). (C) Representative immunostaining images for PECAM-1 and c-Kit. Scale bar is 300 μm. (D) Quantification result of double positive area of PECAM and c-Kit. *p<0.05 versus nondiabetic group (n = 6). (E) Representative immunostaining images of FSP1 and SCF. Scale bar is 300 μm. (F) Quantification result of double positive area of FSP1 and SCF. *p<0.05 versus nondiabetic group (n = 6). (G) Representative immunostaining images of mast cell tryptase and c-Kit. Scale bar is 300 μm. (H) Quantification result of double positive area of Mast cell and c-Kit. *p<0.05 versus nondiabetic group (n = 6).

**Supplemental Figure 7.**
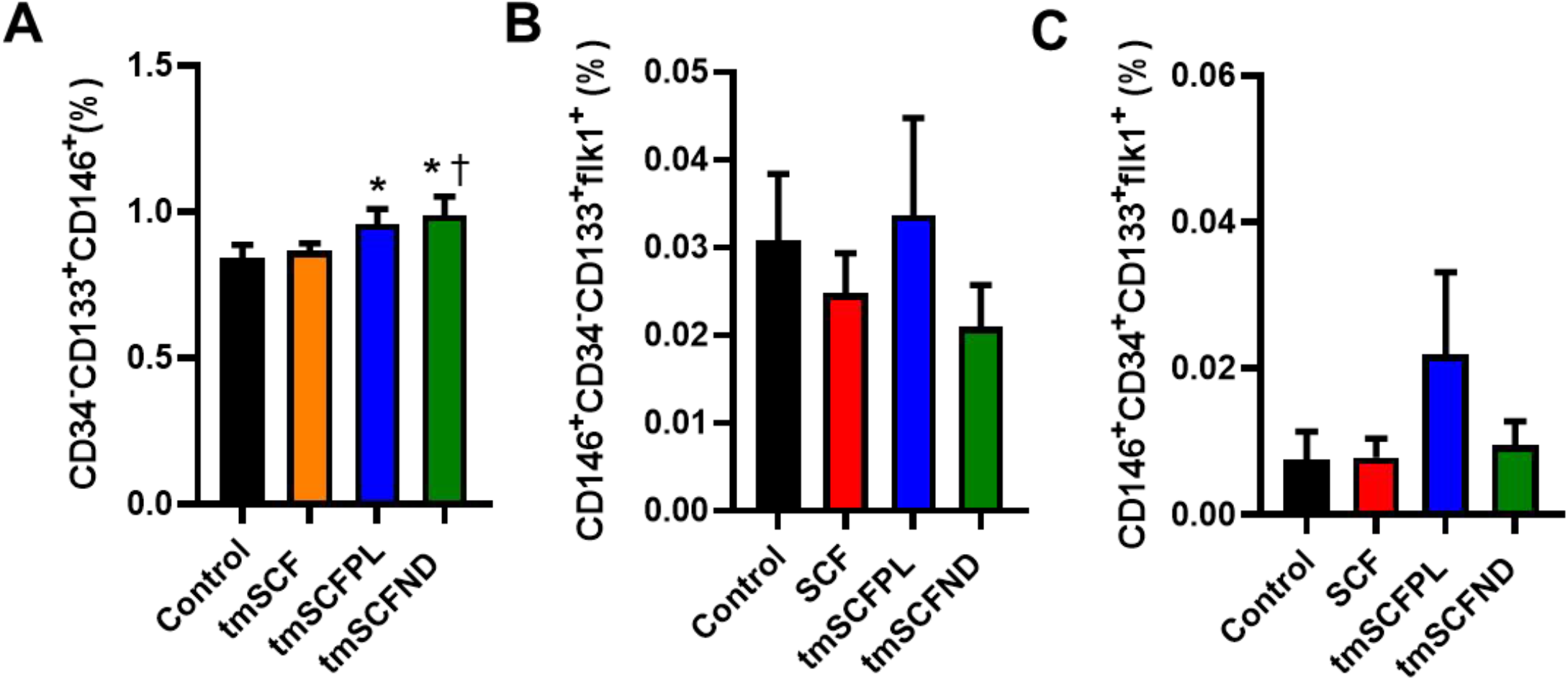
EPC subpopulation was analyzed by flow cytometry. (A) Bone marrow cells treated with our treatments for 30min *in vitro*, then analyzed for CD34^−^CD133^+^CD146^+^ cells population. (B) CD34^−^CD133^+^CD146^+^FLK1^+^ subpopulation in bone marrow was analyzed after four days of subcutaneous injection of our treatment. (C) CD34^+^CD133^+^CD146^+^FLK1^+^ subpopulation in bone marrow was analyzed after four days of subcutaneous injection of our treatment.

## Supplemental Tables

**Supplemental Table 1.** Primary Antibodies Used for Immunostaining.

| Target Protein | Company | Catalog# | Species/Isotype | Dilution Ratio |
| --- | --- | --- | --- | --- |
| GAPDH | Cell Signaling | 21185 | Anti-human | 1:500 |
| PECAM-1 | Cell Signaling | 35285 | Anti-mouse | 1:500 |
| SCF | R&D system | AF-255-NA | Anti-human | 1:500 |
| c-Kit | R&D system | AF1356 | Anti-human, mouse | 1:500 |
| c-Kit | Fisher Scientific | 50-128-29 | Anti-mouse | 1:500 |
| c-Kit | Novas Biological | NBP1-43359 | Anti-mouse | 1:500 |
| FSP1 | Sigma-Aldrich | 07-2274 | Anti-human | 1:500 |
| Mast cell<br>Tryptase | Abcam | Ab2378 | Anti-human | 1:500 |
| Clathrin | Abcam | Ab21679 | Anti-mouse | 1:500 |
| Caveolin | Abcam | Ab2910 | Anti-mouse | 1:500 |
| Phospho-c-Kit | Cell Signaling | 33915 | Anti-mouse | 1:500 |
| αSMA | Abcam | Ab21027 | Anti-mouse | 1:500 |
| CD34 | Santa Cruz | Sc-18917 | Anti-mouse | 1:500 |
| VE-Cadherin | R&D system | AF1002 | Anti-mouse | 1:500 |

